# Awake hippocampal replay is not required for short-term memory

**DOI:** 10.1101/2022.11.03.514989

**Authors:** Lies Deceuninck, Fabian Kloosterman

## Abstract

Short-term memory (STM) on a time scale of seconds to minutes is required to successfully perform day-to-day tasks, for example when engaging in a meaningful conversation. Previous studies in both rodents and primates have correlated hippocampal cellular activity and behavioural expression of STM. This research has led to models describing the putative neural mechanism in the hippocampus that mediate STM. In these models, a key role has been given to hippocampal replay – reactivation of neurons representing a trajectory through space– but definitive causal evidence that can challenge or confirm the model is missing.

In this study, we aimed to address the uncertainty around the role of awake replay in STM by collecting direct causal evidence from behaving rats. Signatures of replay events were detected in the hippocampus and disrupted using electrical stimulation of the ventral hippocampal commissure in rats that were trained on three different spatial memory tasks in a multi-arm radial maze. All tasks required memory of the recent past, but varied in the time scale over which information needed to be retained: (1) a multiple trial match-to sample task, (2) a single trial non-match to sample task and (3) a spatial sequence memory paradigm.

Rats readily learned the task rules, but disruption of awake replay did not affect task performance or other behavioral measures in any of the task. Altogether, our results show for the first time with definitive causal evidence that awake replay is not required for STM of events or of their temporal order.

## Introduction

Short-term memory (STM) is required for many everyday tasks that depend on access to recent events over short time periods, including being engaged in a meaningful conversation and following a set of instructions or previously learned rules. Functional imaging studies (Miyashita and Chang 1988; Ranganath and D’Esposito 2001) and studies with amnesic patients (Hannula, Tranel, and Cohen 2006; Holdstock, Shaw, and Aggleton 1995; Olson et al. 2006) have indicated a role for the medial temporal lobe (MTL) in short-term memory processes, in addition to frontal brain regions (Eichenbaum 2017; Dobbins et al. 2002; Postle 2006; Szczepanski and Knight 2014). However, the neural substrates by which the MTL, and its main component the hippocampus, support short-term maintenance of information are not well understood.

The brain may maintain information in short-term memory through memory rehearsal of recent (task-relevant) experiences (Jonides et al. 2008). It has been hypothesized that at the network level short-lasting neuronal reactivation driven by the hippocampus contributes to memory rehearsal in human (eg. Huang et al. (2018); Tambini and Davachi (2013); Staresina et al. (2013); Liu et al. (2019); Norman et al. (2019)) and rodents (eg. Singer et al. (2013); Pfeiffer and Foster (2013); Xu et al. (2019)) and thus is one candidate mechanism for STM. Hippocampal reactivation has also been linked to learning in general and to memory consolidation, and to date no studies have isolated STM requirements in behavioral tasks to provide causal evidence for the role of hippocampal reactivation in STM.

Of particular interest are hippocampal sequence reactivation (replay) events in the awake state (Foster and Wilson 2006; Diba and Buzsáki 2007). During replay, CA1 place cell firing sequences, that represent a trajectory through space, are re-activated in a time-compressed manner in quiescent episodes immediately after running. These awake replay events may reflect task-relevant past experiences (Foster and Wilson 2006; Karlsson and Frank 2009; Ambrose, Pfeiffer, and Foster 2016; Gupta et al. 2010; H. Freyja Ólafsdóttir et al. 2015; Carey, Tanaka, and Meer 2019) and could be part of a process to temporarily store a memory of that episode into a buffer. Other studies report that replay content seems to relate to the future behavior of animals, potentially guiding behavior, by reflecting trajectories to approach or avoid, or by replaying different navigation options (Singer et al. 2013; Xu et al. 2019; H. Freyja Ólafsdóttir, Carpenter, and Barry 2017; Tang and Jadhav 2018; Shin, Tang, and Jadhav 2019). This activation bias for relevant future trajectories is believed to be a crucial element in current theories for short-term memory, as is the temporary storage buffer (A. D. Baddeley and Hitch 1974; A. Baddeley 2000; RepovŠ and Baddeley 2006). On top of that, when awake replay is disrupted or prolonged during the learning of a short-term memory rule, rats learned the rule significantly slower (Jadhav et al. 2012; Igata, Ikegaya, and Sasaki 2021) or faster (Fernández-Ruiz et al. 2019) respectively. This indirect causal evidence, and the overlap between putative roles of awake replay events reported in the studies above and the expected required processes in theoretical models for short-term memory - the temporary storage buffers and the activation bias - suggests strongly that awake replay is essential for short-term memory.

Due to the lack of direct causal evidence, however, it is still unsure whether the hippocampal replay activity is at the core of short-term memory processes, if it is merely a reflection of short-term memory processes elsewhere in the brain or if it is not involved in short-term memory processes all together. A recent study, for example, showed that replay does not indicate the immediate future action or task relevant information but rather reflects other, more distant past experiences (Gillespie et al. 2021). This observation in combination with the lack of causal evidence leaves room for an alternative hypothesis stating that awake replay events are not at the core of short-term memory processes.

This paper aims to address the uncertainty around the role of awake replay in short-term memory by collecting direct causal evidence from behaving rats. We test the role of awake replay in three different hippocampal dependent behavioral paradigms that each require to use memories of the (recent) past in a different way and over a different time scale; (1) a multiple trial match-to sample task (MTS, (Okaichi and Oshima 1990)), (2) a single trial non-match to sample task (NMTS, (Packard, Hirsh, and White 1989; Sasaki et al. 2018)) and (3) a spatial sequence memory paradigm (SEQ, variation on task described in (Kim and Frank 2009; Jadhav et al. 2012)). These three tasks were chosen because awake replay has been investigated in these tasks directly (Xu et al. 2019) or in variants of these tasks (Jadhav et al. 2012; Fernández-Ruiz et al. 2019)resulting in correlative evidence for awake replay to play a central role in short-term memory processes.

## Results

### Navigation towards recently rewarded locations does not require awake replay

The first short-term memory usage we investigated is the ability to remember and navigate to recently rewarded locations. For this, 5 rats were trained on a spatial match-to sample task (MTS) in an 8-arm radial maze (Figure 1a). In daily training sessions, four randomly selected arms were baited and rats had to learn over the course of 25 trials to only visit these four arms in each trial. As rats learned the reward locations, fewer excursions to non-rewarded arms were made and the number of arm visits per trial decreased (Figure 1b). Rats were trained until the average number of visits per trial for the final 10 trials met the learning criterium consistently for three sessions in a row (Figure 1c).

**Figure 1:**
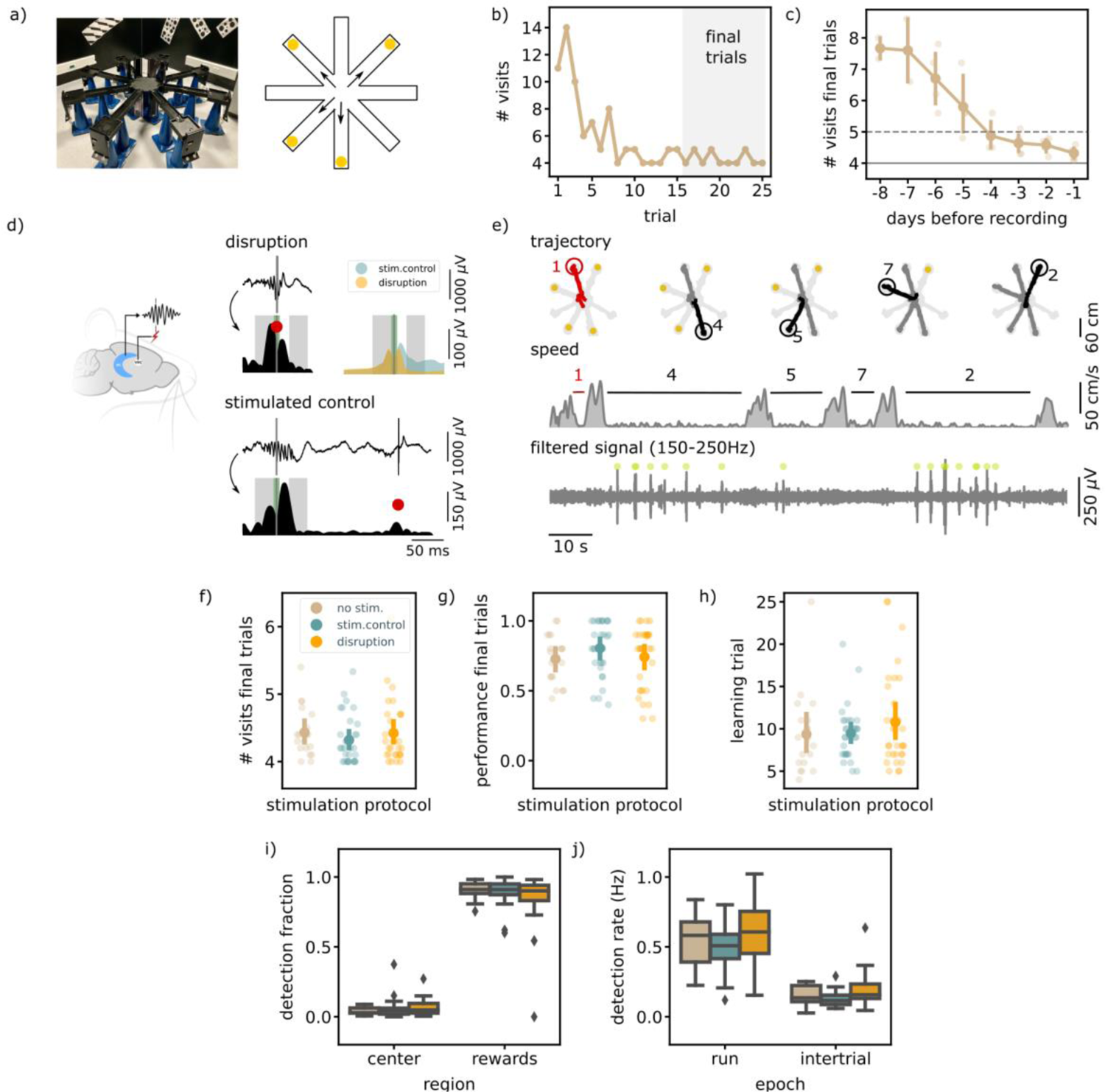
Executing a match to sample task (MTS) is not hippocampal ripple dependent. a) A photo and a schematic illustration of the setup used for the behavior. In a MTS task only four of the eight arms are rewarded. b) Learning curve of one example session showing the total number of visits in each trial. c) The learning curve over pretraining days for all animals (N=5) showing the average number of visits in the final trials per session (small dots; results for individual animals, large dots; mean and 99%CI over animals). The dashed line indicates the learning criteria, the solid line the perfect performance. d) (left) Illustration showing the recording (HC) and disruption (VHC) sites. (right) Example LFP and envelope traces of a disrupted (top) and delayed stimulated (bottom) ripple. The small grey line indicates the time of detection, the red dot the point of stimulation. The green shaded area covers the window in which the stimulation artifact was removed and the grey shaded areas indicate the time windows used to compare the ripple envelope before and after ripple detection. The top right shows the average envelope of all disrupted and delay stimulated ripples in one example session. e) Trajectory, speed and filtered LFP for an example trial. Red trajectory indicates a wrong visit, yellow dot a to be collected reward and the green dot a ripple detection. f) The average number of visits in the final trials per stimulation protocol. g) The average performance (binary quantification) in the final visits per stimulation protocol. h) The learning trial per stimulation protocol. f), g) & h) Small dots represent results for individual sessions (disruption: n= 32, stim.control: n= 31, no stim.: n= 19), large dots reflect the mean and 99%CI per stimulation protocol. i) The fraction of ripple detections during the trials on the rewards and central platforms. The small remaining fraction of detections happened on the arms. j) The ripple detection rate during trials considering only periods of immobility (speed < 5cm/s) and during inter-trial epochs. See also figure 1–figure supplement 1.

Following completion of the training, extracellular field potentials were recorded from hippocampal area CA1 and ripples were detected online as a proxy for sharp-wave ripple (SWR) related hippocampal replay events (Buzsáki 2015; Ciliberti and Kloosterman 2017). In one group of the sessions (N=32) ripple detection was used to trigger closed-loop electrical stimulation of the ventral hippocampal commissure to transiently disrupt ongoing SWR and spiking activity Girardeau et al. (2009) (Figure 1d). The task performance of the rats in these disruption sessions was compared to the performance in the remaining control sessions with either delayed stimulation upon ripple detection (random delay between 150-250ms, N=31) or no stimulation (N=19). Detection and disruption of ripples was performed during all trials and intervening inter-trial intervals. As expected, during trials most ripples occurred during periods of immobility (ripple rate, offline detected: mean [99% CI], 0.52 [0.43,0.61])Hz and reward consumption (Figure 1e). The ripple rate in the inter-trial times is lower than during the trials (mean [99% CI], 0.14 [0.10,0.18]Hz).

The ability of rats to learn the location of the 4 baited arms was not affected by ripple disruption. In the final 10 trials at the end of each session, neither the average number to collect rewards (mean [99% CI]; disruption: 4.42 [4.27,4.70], stim. control: 4.32 [4.18,4.52], no stim.: 4.43 [4.26,4.70], Kruskal-Wallis test: H=2.43, p=0.3, Figure 1f) nor the fraction of trials without any error (disruption: 0.74 [0.63,0.83], stim. control: 0.80 [0.71,0.88], no stim.: 0.73 [0.63,0.83], Kruskal-Wallis test: H=2.56, p=0.28, Figure 1g) differed between disruption and control conditions. To test if the rate of learning over the course of a session was influenced by ripple disruption, we defined a learning trial as the first trial following three correct trials in a row. Rats learned the location of the rewarded arms equally fast in all conditions (disruption: 10.78 [8.75,13.91], stim. control: 9.48 [8.29,11.28], no stim.: 9.37 [7.42,13.81], Kruskal-Wallis test: H=0.70, p=0.71, Figure 1h).

It is possible that the effect of ripple disruption is not reflected in task performance measures, but might be observable in other behavioral parameters. For example, rats may linger on the central platform or arms for longer as an indication of their uncertainty about which arms to visit. For this we first looked at the running speed at the central platform and arms across all three stimulation conditions. Overall, speed was lower on the central platform than on the arms in all three stimulation conditions. When running speed was analyzed separately for arms and center platform, no significant difference was found between the three stimulation protocols (see figure 1–figure supplement 1a). To assess in more detail if rats showed hesitation to perform the task, we divided each session into three stages (early (1-8), middle (9-17) and late (18-25) trials) and looked at the average time spent per arm visit. For all three stages, the average time per visit was not significantly different for disruption condition as compared to the control conditions (Figure 1–figure supplement 1b).

Next, we considered that rats could employ a different strategy in the disruption sessions, which could serve as a compensatory mechanism for the missing awake replay. One well-known strategy for solving a match-to sample task in an 8-arm radial maze is searching for rewarded arms in clockwise or counterclockwise order (Okaichi and Oshima 1990). Rats did not engage in circular choice behavior more often than would be expected by chance for any of the stimulation conditions, and a direct comparison between ripple disruption and control conditions did not reveal a change in circular choice behavior (Figure 1–figure supplement 1c). We noticed that once the rats knew which four arms delivered reward, they tended to visit these arms each time in the same order. To investigate if SWR-associated awake replay influences this stereotypy behavior we computed a stereotypy index for every session by computing the average pairwise similarity between visit sequences of the final trials. The similarity measure was based on the Levenshtein edit distance (Levenshtein 1996) that equals 0 if the visit sequences are the same, and adds 1 for every deletion, insertion or substitution necessary to map one visit sequence onto another. As expected, the stereotypy index was on average higher than expected by chance (disruption: 0.62 [0.55,0.70], stim.control: 0.65 [0.56,0.75], no stim.: 0.59 [0.51,0.67], Figure 1–figure supplement 1c) but was not significantly different between disruption and control sessions (Kruskal-Wallis test: H=0.64, p=0.72).

In all of the above analyses we assumed that each configuration of four arms is equally difficult to learn. To confirm this assumption, we split the configurations in three different categories; (IIa) two sets of neighboring arm pairs, (IIb) one pair of neighboring arms and two arms without direct neighbors and (III) three neighboring arms and one arm without direct neighbors. For this analysis we left out the sessions in which all four arms have no direct neighbor due to a low sample size (n= 2 for each condition). For all three arm configurations, ripple disruption had no influence on any performance quantification, see figure 1–figure supplement 1d.

To look for possible compensatory effects of ripple disruption on network activity, we analyzed the location of ripple detections. In line with literature, the largest proportion of ripples is detected on the reward platforms (disruption: 0.84 [0.68,0.90], stim.control: 0.90 [0.84,0.93], no stim.: 0.90 [0.86,0.93]) followed by the center (disruption: 0.06 [0.04,0.10], stim.control: 0.06 [0.04,0.11], no stim.: 0.04 [0.03,0.06]) and arms (disruption: 0.03 [0.02,0.04], stim.control: 0.02 [0.01,0.03], no stim.: 0.03 [0.02,0.05], Figure 1i). A reward bias parameter for each stimulation protocol, equal to the difference in ripple detections on the rewards and central platform divided by the total number of detections, is not significantly different between the different stimulation protocols (Kruskal-Wallis test: H=1.78, p=0.41), showing that location of the ripple detection did not change across stimulation condition. The online detection rate in the trial (trial) or inter-trial times (inter-trial), calculated using only periods of immobility (<5m/s), was also not different in disruption sessions vs. control sessions (run; disruption: 0.62 [0.52,0.79]Hz, stim.control: 0.50 [0.42,0.56]Hz, no stim.: 0.56 [0.43,0.66]Hz, H=4.71, p=0.095, inter-trial: disruption: 0.19 [0.15,0.26]Hz, stim.control: 0.13 [0.11,0.16]Hz, no stim.: 0.15 [0.11,0.19]Hz, H=5.90, p=0.052, Figure 1j).

Together our results show that SWR-associated awake replay is not required for remembering or navigating to previously rewarded locations over the course of several trials.

### Navigation towards recently non-rewarded locations does not require awake replay

In our second set of experiments we investigated if awake replay might be involved in a different form of short-term memory. One where repetition and thus memory of previous behavior is not enough to solve the task.

Five rats were trained on a non-match to sample task (NMTS) on the 8-arm radial maze, previously described by Sasaki et al. (Sasaki et al. 2018). In every test phase of a trial, rats should visit the four arms that were not rewarded in the preceding instruction phase (Figure 2a, b). Similar to the MTS task most ripples occurred during reward consumption and immobility periods in the instruction and test phases (ripple rate, offline detected, instruction: 0.62 [0.52,0.76]Hz, test: 0.62 [0.52,0.76]Hz). All rats were trained until they reached a learning criterion of 80% fully correct trials in one session for three days in a row (Figure 2c).

**Figure 2:**
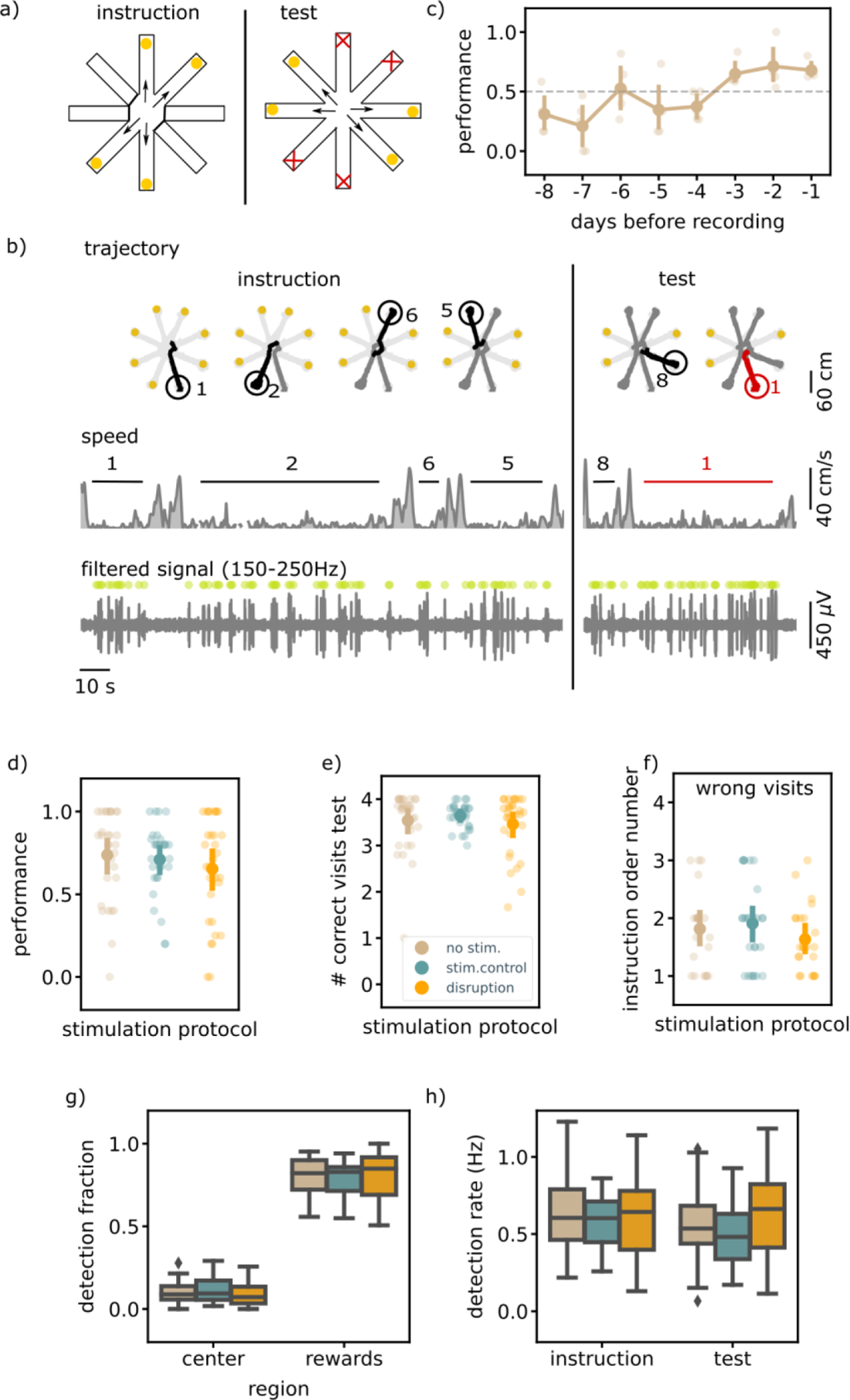
Executing a non-match to sample (NMTS) task is not hippocampal ripple dependent. a) A schematic illustration of the setup used for the behavior. In a NMTS task all eight arms are rewarded but only four can be collected in the instruction phase and the rest needs to be collected in the test phase without an error. b) Trajectory, speed and filtered LFP for an example trial. Red trajectory indicates a wrong visit, yellow dot a to be collected reward and the green dot a ripple detection. c) The learning curve over pretraining days for all animals (N=5) showing the average trial performance in a session. The dashed line indicates the learning criteria (small dots; results for individual animals, large dots; mean and 99%CI). d) The average trial performance in a session per stimulation protocol (no stim: 152, stim. control: 153 and disruption: 147). e) Same as d) but showing the number of visits in the test phase. f) The instruction phase order number of the wrongly visited arm per stimulation protocol. Only wrong trials are considered. (no stim: n= 40, stim. control: n= 42 and disruption: n= 47). For visualization purpose each small dot in d), e) and f) represents a session average. The large dots represent the mean and 99%CI per condition. g) The fraction of ripple detections during the trials on the rewards and central platforms. The small remaining fraction of detections happened on the arms. h) The ripple detection rate during the instruction and test phase of the trials, considering only periods of immobility (speed < 5cm/s). See also figure 2–figure supplement 1.

Once the learning criterion was met, ripple disruption was performed in one third of the trials in each session, with the remaining trials constituting delay stimulation and no stimulation control trials. Ripple detection and disruption always happened during all phases of the trial (instruction and test). The trial performance (i.e., whether or not a trial is completed without a single error) and number of correct visits in the test phase in the disruption trials were not significantly different from those in the control trials (performance; disruption: 0.68 [0.57,0.77], stim.control: 0.73 [0.62,0.80], control: 0.74 [0.63,0.82], Kruskal-Wallis test: H=1.30, p=0.52, correct visits test phase; disruption: 3.52 [3.29,3.67], stim.control: 3.65 [3.48,3.76], control: 3.57 [3.32,3.72], H=1.44, p=0.49, Figure 2d & 2e).

For rats to correctly perform the task and identify which arms have not yet been visited, one strategy is to memorize all four visits from the instruction phase. We observed a temporal order effect such that errors in the test phase are biased to the arms that were instructed furthest back in time (average order number instruction arm; disruption: 1.70 [1.43,2.01], stim.control: 1.93 [1.60,2.26], no stim.: 1.88 [1.55,2.25], Figure 2–figure supplement 1c). We observed no difference between trials with or without ripple disruption (Kruskal-Wallis test: H=1.79, p=0.41, Figure 2f), which taken together with the previous quantifications strongly suggests that the short-term memory for recently non-rewarded locations is not impaired by SWR-associated replay disruption.

We quantified other behavioral parameters to investigate if replay disruption has a significant impact there. Similar to the results in the MTS task, rats ran faster on the arms than on the central platform (Figure 2–figure supplement 1a) suggesting that rats take time on the central platform to select the next arm to visit. If awake replay is important for identifying the remaining baited arms, rats might be more uncertain about their choice in disruption trials which could be reflected in more lingering at the central platform. We observed no difference in speed between trials in a different stimulation protocols both on the central platform and arms (Figure 2–figure supplement 1a). Likewise, the time per visit in the test phase was not impacted by ripple disruption (Figure 2–figure supplement 1b).

Due to a difference in experimental setup (see methods), the inter-phase time for three animals was longer and more varied compared to the inter-phase times for the two other animals (Figure 2–figure supplement 1e). To test if trials with longer intervals may be more susceptible to ripple disruption, we compared the task performance between both groups of animals. We observed a tendency for rats to perform better with short inter-phase intervals, but there was no difference between stimulation conditions for either short or long intervals (Figure 2–figure supplement 1f).

Similar to the MTS task, the largest fraction of ripples were detected when the rats were on the reward platform (rewards; disruption: 0.81 [0.74,0.86], stim.control: 0.78 [0.72,0.83], no stim.: 0.80 [0.74,0.84], center; disruption: 0.09 [0.06,0.13], stim.control: 0.11 [0.08,0.15], no stim.: 0.10 [0.07,0.13], arms; disruption: 0.05 [0.03,0.08], stim.control: 0.06 [0.04,0.08], no stim.: 0.06 [0.04,0.08], Figure 2g). The spatial distribution of ripple occurrence was not influenced by ripple disruption overall (Kruskal-Wallis test: H=1.56, p=0.46) nor in either instruction or test phase (Figure 2–figure supplement 1d). The online ripple detection rate, calculated using only periods of immobility (<5m/s) in the trials was also not significantly different between trials with a different stimulation protocol in both the instruction and test phase (instruction: disruption: 0.61 [0.49,0.73]Hz, stim.control: 0.58 [0.50,0.65]Hz, no stim.: 0.64 [0.54,0.76]Hz, Kruskal-Wallis test: H=0.73, p=0.7, test: disruption: 0.63 [0.50,0.75]Hz, stim.control: 0.49 [0.41,0.58]Hz, no stim.: 0.55 [0.44,0.67]Hz, Kruskal-Wallis test: H=4.29, p=0.12, Figure 2h).

### Navigation using temporally ordered information does not require awake replay

The results from the first two experiments indicate that awake replay is not involved in the short-term memory of recent events. It begs the question what aspect of *learning a short-term memory rule* is supported by awake replay (Jadhav et al. 2012; Fernández-Ruiz et al. 2019).

The short-term memory rule that rats were unable to learn due to ripple disruption in these studies, required navigation from a center arm to two outer arms in an alternating fashion; left, center, right, center, left,… To learn this rule, it is crucial to remember the previous events in the correct temporal order, something that was not required in either the MTS or NMTS task.

To test if hippocampal replay is required for memorizing the temporal order of two or more recent events, we designed a spatial sequence memory paradigm (Figure 3a). In this task, rats need to learn each day the order in which to visit a set of four new arms by remembering which transitions are rewarded. After extensive pre-training on the linear track and three-arm radial maze, five rats were pre-trained on the sequence memory paradigm for several days until the final average sequence performance (average sequence performance in the last 100 trials) was above 50% for at least two sessions in a row. The sequence performance is calculated by multiplying the outcome of all individual visits (1= correct transition, 0= incorrect transition) in a moving window of five visits. A correct sequence performance, that means five correct visits in a row, will yield a sequence performance of one (Figure 3 b, c). Ripples are mostly detected on the reward platform when the animal is immobile (ripple rate, offline detected, 0.45 [0.40,0.51]Hz, Figure 3d). The ripple rate during the interleaving rest epoch is comparably high (ripple rate, offline detected, 0.54 [0.50,0.59]Hz).

**Figure 3:**
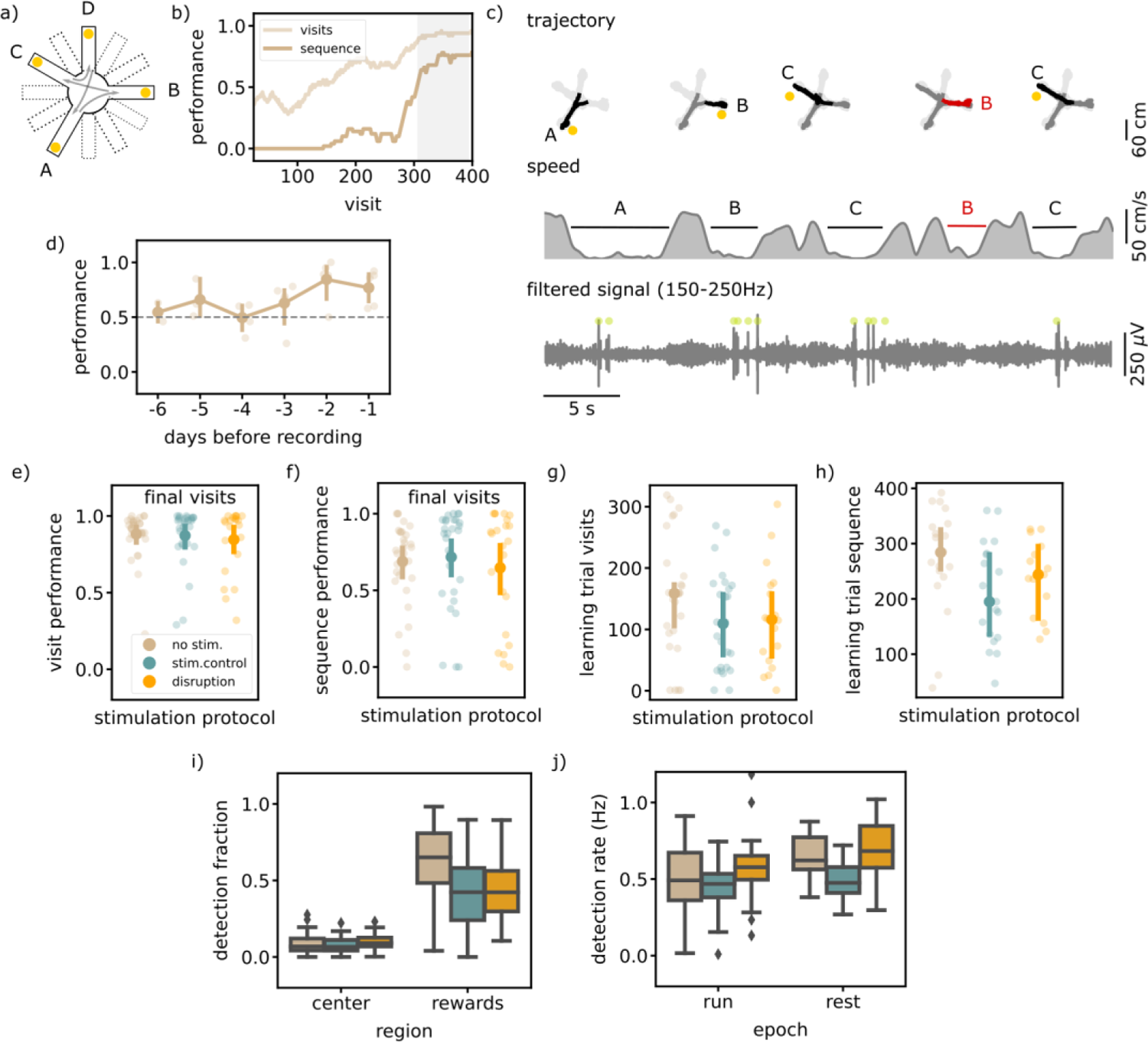
Executing a sequential (SEQ) spatial memory task is not hippocampal ripple dependent. a) A schematic illustration of the setup used for the behavior. In a SEQ task only four of twelve possible arms are present. Reward is delivered when arms are visited in the correct order. b) Learning curves of one example session showing the visit performance and sequence performance over visits, both visualized using a moving average with N=50. c) Trajectory, speed and filtered LFP for five visits in an example session. Red trajectory indicates a wrong visit, yellow dot a to be collected reward and the green dot a ripple detection (small dots; results for individual animals, large dots; average over animals). d) The learning curve over pretraining days for all animals (N=5) showing the average sequence performance in the final 100 visits in a session. The dashed line indicates the learning criteria (small dots; result per animal, large dots: mean and 99%CI). e) The average visit performance over the final visits of a session per stimulation protocol. One small dot represents one session (no stim: 31, stim. control: 29 and disruption: 23) and the large dot the mean and 99%CI. f) Same as e) but showing the average sequence performance over the final visits of a session. g) The visit learning trial of sessions per stimulation protocol defined using the smoothed learning curve of the visit performance(see methods). Only session where learning criteria was met within 400 visits are included. h) Same as g) but showing the sequence learning trial computed using the smoothed learning curve of the sequence performance. i) The fraction of ripple detections during the trials on the rewards and central platforms. The small remaining fraction of detections happened on the arms. j) The ripple detection rate during run on the maze, considering only periods of immobility (speed < 5cm/s), and during rest epochs in the sleep box. See also figure 3–figure supplement 1.

To test the involvement of awake replay in memorizing and learning a spatial sequence we disrupted ripples in one third of the sessions (N = 23), throughout all sessions each day; the two run sessions and rest session. The other sessions were used for stimulated (N = 29) and no-stimulated (N = 31) controls, similarly as defined for the MTS and NMTS task.

The average visit and sequence performance in the final visits indicated no significant difference between the different stimulation protocols (visit performance; disruption: 0.85 [0.71,0.93], stim.control: 0.87 [0.74,0.94], no stim.: 0.88 [0.77,0.93], Kruskal-Wallis test: H=0.75, p=0.69, sequence performance; disruption: 0.65 [0.42,0.81], stim.control: 0.72 [0.53,0.84], no stim.: 0.69 [0.55,0.79], Kruskal-Wallis test: H=1.03, p=0.6, Figure 3e, f). To test if rats learn at a different rate, we computed for every session the smoothed learning curve of both the visit and sequence performance and defined the visit where the curve is significantly above 0.5 as the learning visit. In every stimulation condition there was a similar percentage of sessions in which the rat does not reach the learning criteria, which we left out for this quantification (visit performance; disruption: n= 5, stim. control: n= 3, no stim: n= 4, sequence performance; disruption: n= 8, stim. control: n= 10, no stim: n= 11). The learning visit for both the visit and sequence performance was not significantly different between sessions with a different stimulation protocol (learning visit visit performance; disruption: 117.72 [73.47,172.90], stim.control: 112.69 [77.20,154.96], no stim.: 154.22 [108.18,201.02], Kruskal-Wallis test: H=2.66, p=0.26, learning visit sequence performance; disruption: 238.80 [193.51,280.09], stim.control: 209.21 [154.85,261.91], no stim.: 273.10 [202.75,316.72], Kruskal-Wallis test: H=6.01, p=0.05, Figure 3g,h).

The overall behavior quantifications also did not show any change due to ripple disruption. Like in the previous experiments we quantified the speed on the central platform and arms and looked at the time per visit as a measure for how certain rats were about their choice. The speed on the central platform and arms showed no difference in different stimulation conditions (Figure 3–figure supplement 1a). Rats also did not take more or less time per visit in disruption vs. control trials (Figure 3–figure supplement 1b).

When we looked at the smoothed learning curves for the individual transitions, we observed that not all transitions were learned equally fast (Figure 3–figure supplement 1c). This is an indication that rats are probably not learning individual transitions, but are aware that there is a repeating pattern to be found. Namely, the possibilities for the new transitions are lower when using the information of learned transitions in the correct temporal order. By looking at the learning visits for the individual transition learning curves, we can investigate if the strategy employed by the rats changes due to ripple disruption. For all three stimulation protocols, learning visits of individual transitions vary and the distributions were significantly different between stimulation protocol (Figure 3– figure supplement 1d). Post-hoc Mann-Whitney tests however revealed only a significant difference between the two control conditions (Figure 3–figure supplement 1d). Next, we quantified the difference between the learning visits of the slowest and fastest learned transition in sessions where the rat had learned at least two transitions, as a measure of how much use had been made of the repeating pattern. This quantification of the learning trial difference in each session showed no significant difference due to ripple disruption (Figure 3–figure supplement 1e).

In line with our expectations, ripples were detected mostly on reward sites (rewards; disruption: 0.44 [0.37,0.51], stim.control: 0.42 [0.34,0.49], no stim.: 0.62 [0.55,0.69], arms; disruption: 0.05 [0.03,0.06], stim. control: 0.03 [0.02,0.05], no stim.: 0.04 [0.03,0.05], center; disruption: 0.10 [0.08,0.12], stim. control: 0.08 [0.06,0.09], no stim.: 0.08 [0.07,0.11], Figure 3i), and this bias was not significantly different between stimulation protocols (Kruskal-Wallis test: H=1.60, p=0.45). The overall ripple detection rate calculated over the immobility periods in both run epochs and in the rest epochs did seem to be significantly different between sessions with a different stimulation protocol (run; disruption: 0.58 [0.52,0.68], stim.control: 0.46 [0.41,0.50]), no stim.: 0.49 [0.41,0.57], Kruskal-Wallis test: H=12.94, p=0.0016, rest; disruption: 0.69 [0.58,0.79], stim. control: 0.49 [0.43,0.54]), no stim.: 0.64 [0.58,0.70], Kruskal-Wallis test: H=21.02, p=2.7 × 10^−5^). However, post-hoc Mann-Whitney tests for the run epoch revealed no significant difference between any pair (disruption - stim.control: U=1944.00, p=6.6 × 10^−5^, disruption - no stim.: U=1728.00, p=0.061, stim.control - no stim.: U=2011.00, p=0.26). In the rest epoch the post-hoc Mann-Whitney tests revealed a significant difference between the disruption and stimulated control sessions but not between the other pairs (disruption - stim.control: U=520.00, p=0.00019, disruption - no stim.: U=435.00, p=0.17, stim.control - no stim.: U=695.00, p=7.7 × 10^−5^). Together these quantifications suggest that ripple disruption has no overall effect on the hippocampal activity.

### Awake hippocampal ripples are accurately detected and disrupted

Before each experiment, the threshold for ripple detection and the stimulation current were manually adjusted by the experimenter to ensure accurate and complete detection and disruption in every session. Post-hoc quality check of both the disruption and detection was done in the same way as Michon et al. (Michon et al. 2019). The disruption quality was quantified for all session in stimulated control and disruption condition by comparing the amplitude in a 20ms window before and after the ripple detection. The normalized post-detection ripple amplitude was significantly lower in the sessions with ripple disruption in all three tasks (Mann-Whitney test, MTS: U=961.00, p=7 × 10^−12^, NMTS: U=912.00, p=5.9 × 10^−11^, SEQ: U=2569.00, p=3.6 × 10^−18^), showing that all detected ripples were disrupted (Figure 4a). The quality of the ripple detection was quantified using the stimulated and non-stimulated control sessions. In each session ripples were detected off-line (see methods) and by comparing these with the on-line detected ripples both the true positive rate (TPR) and false discovery rate (FDR) were quantified (Figure 4b). The high TPR for almost all sessions suggest that a large fraction of all ripples were detected. We observed a wider distribution for the FDR in comparison to the distribution for the TPR, likely due to a variance in signal to noise ratio per experiment. Ripple detection and disruption were quantified separately for the inter-trial times in the MTS task and the rest epochs in the SEQ task (Figure 4–figure supplement 1). In both cases, ripple power was significantly reduced following stimulation (Figure 4–figure supplement 1a). The TPR-FDR distribution for the inter-trial times in the REF task was shifted towards lower TPR and higher FDR, indicating a worse detection (lower TPR and higher FDR, Figure 4–figure supplement 1b). In the inter-trial times the animals were locked in the center of the maze and they frequently engaged with the closed doors, which resulted in movement artifacts in the LFP signal that were falsely identified as ripples. Quantifications for the sleep box epochs in the SEQ task were in similar to those on the maze, that is high TPR and spread out FDR (Figure 4–figure supplement 1c). Because awake ripple first need to occur before we can detect them, there will always be a detection delay (Figure 4–figure supplement 1c). The absolute detection delay over all epochs in each task, and considering only the no stim. and stim.control epochs, is median [iqr], 36.69 [34.68,39.44], in line with what we expect when using falcon (Ciliberti and Kloosterman 2017). When we compare this to the total length of the ripple we obtain an relative delay of median [iqr], 52.90 [49.62,55.80].

**Figure 4:**
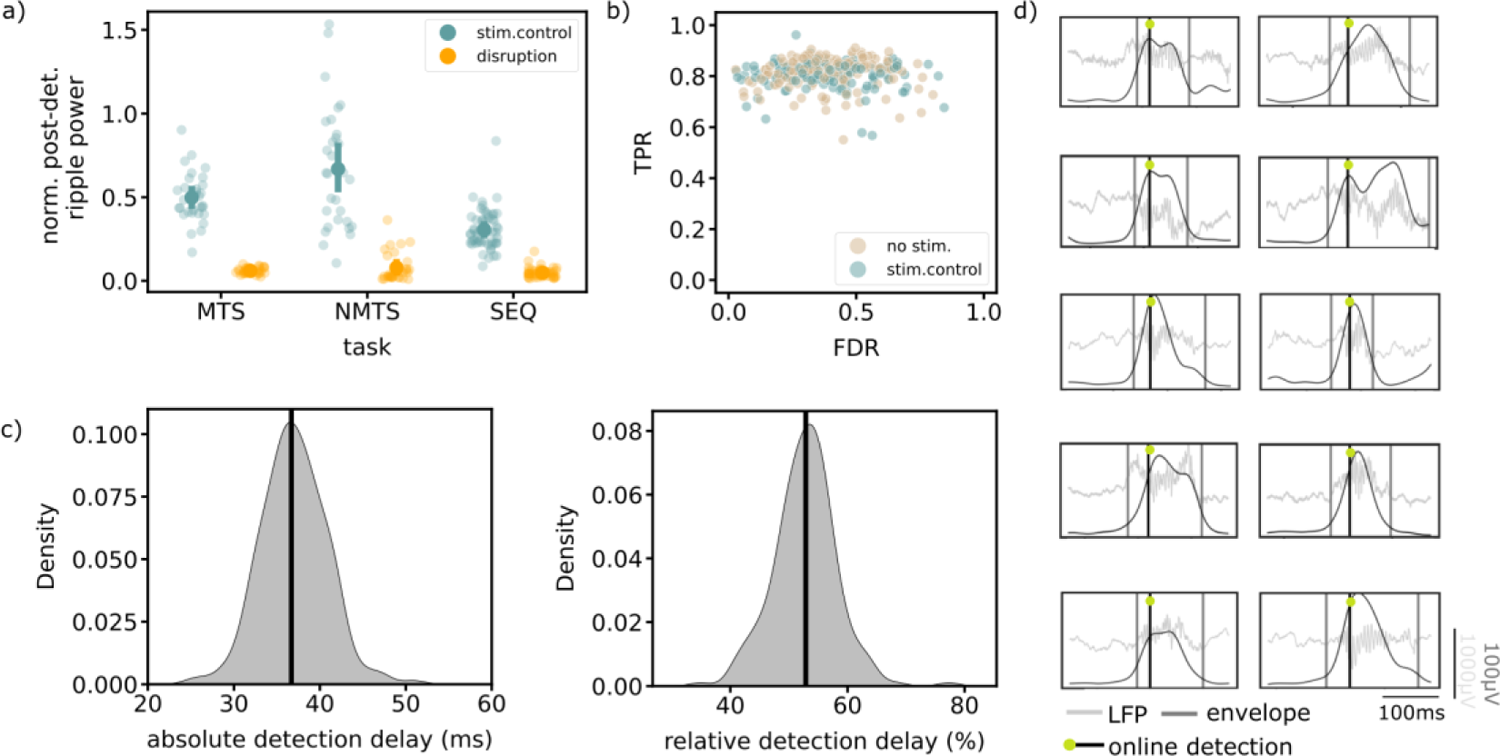
Available quantifications indicate accurate ripple detection and disruption. a) Normalized post-detection ripple power for all tree tasks in run epochs. A small dot represents the average over one session, the large dots represent the mean and 99%CI. b) True positive rate versus False discovery rate for all tasks. One dot represents the average over one session. c) The absolute (left) and relative (right) detection delays for all epochs of all tasks, considering only the no stim. and stim.control epochs. d) Example online detected ripples that were also detected offline (true positive) from five different animals. Grey vertical lines indicate the start and end of the ripple. See also Figure 4–figure supplement 1.

## Discussion

Altogether, our results show for the first time with direct causal evidence that awake replay is not involved in short-term memory of events or the temporal order in which they happened. This conclusion is drawn from three different sets of experiments, where animals were well accustomed to both the mazes and task rules, excluding any interference of the learning and consolidation of rules and environment representations. Our results also revealed that replay disruption had no influence on other behavioral parameters or on the frequency and location of the replays. Overall these results suggest that awake hippocampal replays are not mediating the storage of information to temporary storage buffers nor do they partake in the process to make appropriate navigation decisions using short-term memories.

Awake replay has been analyzed in very similar match-to-sample and non-match-to-sample paradigms (Xu et al. 2019; Sasaki et al. 2018), both shown to be hippocampal dependent (Okaichi and Oshima 1990; Packard, Hirsh, and White 1989). Replay trajectories are compared to the task rules and navigation choices of the animal, and even though correlations were found, these sometimes do not agree with the working model or leave room for other interpretations. For example, Xu et al. (2019) observed that in a match to sample task replay relates to navigation choices only when a correct choice is made, not when incorrect choices are made. If replay is truly guiding navigation, it should predict both correct and incorrect choices. In other cases like for Ambrose, Pfeiffer, and Foster (2016) or Carey, Tanaka, and Meer (2019) it is difficult to make a distinction between a replay that correlates to a next choice or a replay of a past experience due to a limited number of navigational choices. The recent study by Gillespie et al. (2021) tackled this problem by designing a new short-term memory paradigm that allows the clear distinction between replay of future and past paths by having many spatially and temporally distinct navigational options. They interestingly observed no correlation between replay content and navigation choices. Rather the replay seemed to represent seemingly non-relevant past trajectories, arguing against a role for awake replay in short-term memory. Our experiments provide causal evidence for this hypothesis.

A number of studies suggest that the hippocampus is important to remember the temporal order of events (Gilbert and Kesner 2002; Fortin, Agster, and Eichenbaum 2002; Kesner, Hunsaker, and Gilbert 2005; Farovik, Dupont, and Eichenbaum 2010). To test the role of awake replay in this brain function, we modified the known hippocampal dependent 3-arm spatial alternation paradigm (Kim and Frank 2009) into a sequence memory paradigm that requires the rat to learn to visit four location in a specific temporal order. A better temporally ordered memory of previously visited arms helps to learn this order faster. In this third experiment (SEQ task) ripple disruption did not interfere with the speed with which animals learn the spatial order showing that awake replay does not support remembering the temporal order of events. Future research will have to point which other hippocampal activity supports sequence memory. Yet, we acknowledge that despite the high similarity with known hippocampal dependent spatial tasks, we cannot exclude that the lack of involvement of awake replay is merely a consequence of the fact that this new paradigm is not hippocampal dependent.

Zhang et al. (2021) and Jarzebowski et al. (2021) inactivated medial septum neurons during the execution of a learned short-term memory task. The resulting performance deficit correlated with a decrease in SWR occurrence, which suggested a crucial role for awake replay in the task execution. Our results presented here use a direct manipulation of hippocampal replay activity and show no influence on the short-term memory performance. The suppression of SWRs observed by Zhang et al. (2021) and Jarzebowski et al. (2021), as a result of increased activity of cholinergic neurons by optogenetic stimulation, are caused by hitherto unidentified mechanisms, independent of the SPW-R-suppressing effect.

We observed a clear drop in ripple amplitude after each VHC stimulation, strongly indicating that detected ripples are successfully disrupted. Due to the filtering of the LFP signal, which is necessary for the identification of ripples, ripples are detected with a short latency. It is unlikely however that this would be the reason for an absence of deleterious effect on the short-term memory performance as ripple disruptions with similar latencies caused behavioral effects in previous experiments (Michon et al. 2019). Moreover, long ripples (>120ms) contribute most strongly to memory processes (Fernández-Ruiz et al. 2019) and with our detection delay of median [iqr], 36.69 [34.68,39.44] are disrupted for more then 70%.

When comparing the behavioral outcomes across studies in which ripple events are disrupted, it is important to check that the manipulations were performed in a similar manner. Which ripples are detected is dependent on the employed real-time algorithm, on the signal to noise ratio of the recorded signals and on the used detection threshold. Setting the threshold too high minimizes the false discovery rate, but with the risk of missing a large fraction of small ripples. The ripple detection rates in our experiments are in line with what is reported in literature, which increases our confidence that ripples were detected accurately and in a similar way as before. In addition, comparison of the online detections with an offline detection algorithm yielded a high true positive rate, meaning that almost all ripples detected offline are also detected online.

We acknowledge that our study does not include a behavioral task that was positively affected by ripple disruption (i.e., a “positive control”) and could provide further demonstration that our implementation of the disruption approach is sound. However, previous work from our lab employing the same method for disruption of sleep ripples showed a clear behavioral effect (Michon et al. 2019). In addition, the size of reduction in ripple power after the electrical stimulation in the present study is comparable to studies from other labs that used the same approach (Michon et al. 2019; Girardeau et al. 2009; Jadhav et al. 2012; Fernández-Ruiz et al. 2019). Finally, power analysis shows that the number of sessions and trials we have in all three experiments should allow us to pick up with 95% confidence at least a 10% mean difference with 80% power, which is in line with previous SWR disruption experiments with observed performance differences in the range of 10-20% (Michon et al. 2019; Girardeau et al. 2009; Jadhav et al. 2012; Fernández-Ruiz et al. 2019). Overall, these considerations increase our confidence that if awake replay were indeed important for successful performance of any of the three tasks, we would have observed so.

In summary our results provide evidence that awake replay is not involved in memorizing past events and their temporal order over a short period of time. Several studies show that awake replay is required for the acquisition of a task rule that requires short-term memory (Jadhav et al. 2012; Igata, Ikegaya, and Sasaki 2021; Fernández-Ruiz et al. 2019). By extension, several computational models that have virtual agents learn a similar foraging task have significantly increased the learning performance by incorporating replay like activity (Mattar and Daw 2018; Hayes et al. 2021; Roscow et al. 2021; Johnson and Redish 2005). Learning a new task rule, however, is a complex iterative behavior that requires different brain functions, such as information registration, pattern inference, revision and goal-directed modulations, to coordinate together at different time scales. To find out what role awake replay plays in learning, specific causal experiments need to be designed that can isolate and investigate the different aspects of learning separately.

## Acknowledgments

We thank Jyh-Jang Sun, Frédéric Michon and Ta-Shun Su for help with the development of the sequence paradigm and Marine Chaput for setting up the automated maze setup and live video tracking. L.D is funded by Research Foundation Flanders (FWO), Belgium as a PhD Fellow fundamental research, grant number 11D9322N. F.K. is funded by Research Foundation Flanders (FWO),Belgium under grant number G077321N and KU Leuven, Belgium C1 grant C14/17/042

## Author Contributions

Conceptualization, L.D and F.K.; Methodology, L.D and F.K.; Software, F.K.; Formal Analysis, L.D.; Investigation, L.D.; Writing – Original Draft, L.D; Writing – Review & Editing, L.D and F.K.; Visualization, L.D; Supervision,F.K.; Funding Acquisition, L.D. and F.K.

## Methods

### Experimental model and subject details

A total of 15 male Long Evans rats (supplier Janvier Labs), food restricted to 85%–90% of the free-feeding weight, were used in this study. At the start of behavioral training procedures, rats were 7-10 weeks old. All rats received an implant for neural recordings and stimulation. Tables 1-3 provide an overview of the experimental sessions for each rat. All experiments were carried out in accordance with protocols approved by KU Leuven animal ethics committee (P119/2015 and P175/2020) and in accordance with the European Council Directive, 2010/63/EU. Animals in experiments were housed separately in individually ventilated cages (IVC) with ad libitum access to water and controlled intake of standard food pellets. Health status and body weight were checked daily by the experimenters and dedicated animal care personnel. To improve the welbeing of the rats, a playpen (100 x 55 cm) was constructed in the course of this study that allowed rats to spend spent time in an enriched environment with their (former) cage mate. The playpen can be divided into two parts separated by a transparent wall with ‘sniffle holes’ to encourage social interaction between the two rats without accidental damage to their implants. The enrichment consisted of a small maze with hidden food, toys, a climbing rope and ladder that provide access to an upper story with a running wheel. Experiments for the delayed non-match to sample and match to sample tasks were started prior to completion of the playpen. All five rats used to test the memory of temporally ordered information (third experiment), however, spent at least 30 min per day in the playpen.

**Table 1:** Overview animals MTS

| **Animals** | **Disruption sessions** | **Stim.control sessions** | **No stim sessions** |
| --- | --- | --- | --- |
| LD10 | 4 | 3 | 1 |
| LD11 | 3 | 2 | 3 |
| LD14 | 9 | 10 | 6 |
| LD21 | 8 | 8 | 4 |
| LD22 | 7 | 8 | 3 |
| **Total** | **31** | **31** | **17** |

**Table 2:**
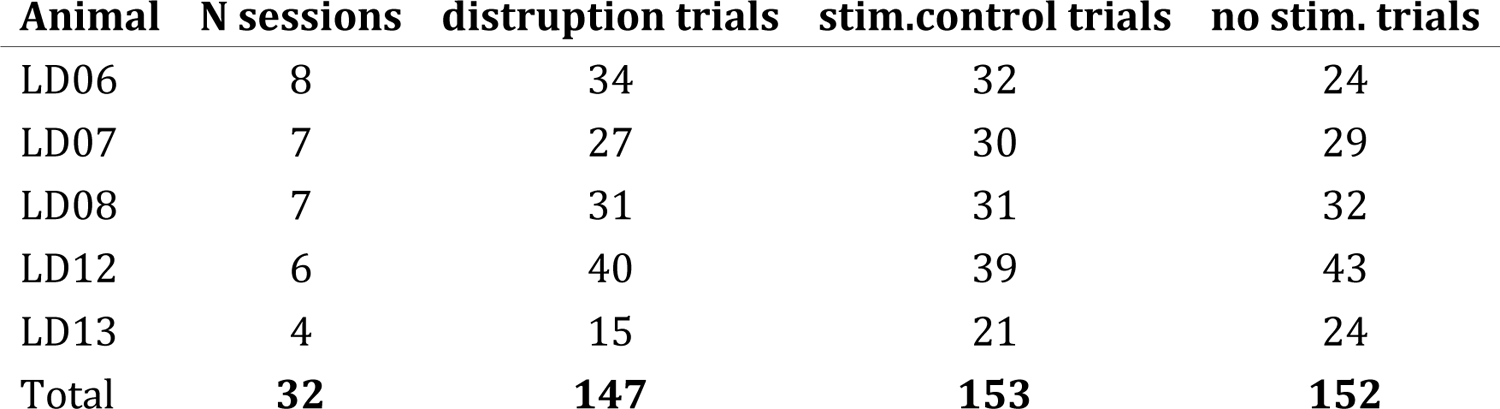
Overview animals NMTS

**Table 3:** Overview animals SEQ

| **Animals** | **Disruption sessions** | **Stim. Control sessions** | **No stim. sessions** |
| --- | --- | --- | --- |
| LD53 | 7 | 7 | 7 |
| LD56 | 8 | 7 | 6 |
| LD57 | 5 | 6 | 6 |
| LD58 | 5 | 5 | 6 |
| LD60 | 6 | 4 | 4 |
| **Total** | **31** | **29** | **29** |

### Behavioral task

In all experiments, the maze is elevated 40 cm above the ground and located in a 4 x 4 m black room with distinctive visual cues on all four walls.

#### Match-to-sample paradigm (MTS)

We use an 8-arm radial maze with a 40 cm diameter central platform and 90 cm long arms that each terminate in a 20 cm squared reward platform. There are doors at 6 cm from the central platform on each arm, which open by moving down and are controlled wirelessly from outside the room. Three animals received chocolate pellets as reward, delivered in a small food well at the end of the reward platform. The remaining two animals received a liquid reward (chocolate syrup), dispensed through a pump that was also controlled wirelessly from outside the behavior room. The goal of the task is for rats to learn in daily sessions which set of four out of eight arms lead to reward. Each session consists of 25 trials and the four rewarded arms are randomly varied between sessions, such that each session is an independent assessment of the rats’ ability to learn the location of reward. Configurations that have all four arms located next to each other were not used. When a solid reward is used, a trial starts with the four chosen arms baited. The other arms contain inaccessible rewards to minimize olfactory strategies. When liquid rewards are used, a pump containing chocolate syrup is placed on every arm to minimize olfactory and visual strategies. Only the pumps on the correct arms deliver reward (∼2ml) when the rat visits that arm. At the start of a session, rats are placed on the central platform, the experimenter leaves the room, and all doors open simultaneously. Rats can now explore all arms freely. After the last rewarded arm has been visited, the rats are allowed to return to the central platform, after which all doors close and the first trial ends. In case solid rewards are used, the reward wells of the same four arms are now refilled by the experimenter. Rats remain on the central platform for 60-120s, after which all doors are opened and the next trial starts.

#### Non-match to sample paradigm (NMTS)

We use the same apparatus as for the MTS task with the doors controlled wirelessly for all animals. All animals received chocolate pellets as a reward, delivered in a small food well at the end of the reward platform. The doors are controlled wirelessly in sessions of two animals, for the sessions of the other three the doors are controlled manually. The goal of this task is for rats to remember which arms in an 8-arm radial maze they had and had not visited recently (Sasaki et al. (2018)). In each daily session, rats perform 15 independent trials. Every trial consists of two phases: an instruction phase in which rats are forced to visit 4 randomly selected arms one by one, and test phase in which rats have access to all 8 arms and need to visit the remaining 4 arms to collect reward (Figure 2a). At the start of each trial, all eight arms are baited with a chocolate reward at the end of the reward platform. The instruction phase starts with the rat in the center of the maze with all doors closed. Next, one after the other, 4 randomly chosen doors are opened and the rat is allowed to run down the arm and collect the reward. After each arm visit the corresponding door closes so that at any moment only one arm can be visited. After the fourth visit all doors are closed, the instruction phase ends and the rat is constrained to the central platform, in case automated doors (Peira bvba) were used, or placed on a nearby platform next to the maze, when manual doors were used. After a short delay all doors open, rats are returned to the central platform if needed, and the test phase started. After a correct visit in the test phase, the door of that arm was closed for the remainder of the trial. A trial ended after rats visited all four correct arms or after the first incorrect visit. In between trials, rats either waited for 90-120s in the center of the maze with all doors closed (in case of automated doors) or were placed on a nearby platform (in case of manually operated doors). During the inter-trial wait period, all reward wells were refilled by the experimenter.

#### Sequence memory paradigm (SEQ)

For the sequence memory paradigm we use a 12-arm radial maze with a 37.5 cm diameter central platform and 60 cm long arms that each terminate in a 22 cm squared reward platform. There are no doors present and all animals receive a liquid reward (condensed milk, ∼2 ml) that was delivered wirelessly from outside the behavior room. In the sequence memory task the goal is for rats to learn the correct order in which four arms need to be visited to receive reward. Each daily session is a separate experiment in which four arms of a 12-arm radial maze are randomly chosen, and the remaining arms are removed. A crossing visit order is selected (i.e., clockwise and counterclockwise orders are not used), for example if arms in positions 2,3,4,11 are chosen then valid visit orders are for example 2→4→3→11→2 and 2→3→11→4→2, but not the counterclockwise order 2→11→4→3→2. Every session consists of two run epochs lasting 25 min and an intervening rest epoch lasting 10 min. At the beginning of the session rats are placed in the center of the maze with all doors closed. The first run epoch starts when all doors open and the rat is free to move around. Each time a correct transition is made to the next arm in the sequence the rat is rewarded. After 25 min rats are allowed to return to the center, the doors are closed and rats are taken off by the experimenter and put back in the sleep box for the rest epoch. The second run epoch is identical to the first. The rats’ position is tracked by an overhead camera and liquid rewards are automatically delivered (Peira bvba) based on the tracked behavior using custom developed software in the lab.

### Behavioral training

All animals were pretrained to run on elevated mazes for reward, habituated to automated doors (when used) and to a separate sleep box. For each of the three behavioral tasks, the pretraining and training procedures to teach the rats the task rules are described below.

#### Match-to-sample paradigm

In the first phase of pretraining (5-7 days), rats are habituated to the 8-arm radial maze and learn to look for reward at the end of the arms. On days 1-2, rats are placed in the center of the maze and allowed to freely explore for 15-30 min. Chocolate rewards are scattered throughout the maze to promote exploration. On the next 2-3 days, rewards are progressively restricted to only the arms and finally the reward wells at the end of each arm. Once rats are sufficiently comfortable and motivated to run for the rewards, the second phase of the pretraining starts. In this phase, rats perform trials of the MTS paradigm until they complete at least 15 trials in 30 minutes. Training is completed after recovery from surgery when rats are back at 85-90% of their post-surgery weight. In daily sessions (25 trials), rats perform the MTS paradigm until the average number of visits on the last 10 trials is less than 5 for three days in a row.

#### Non-match-to-sample paradigm

The pretraining prior to surgery is the same as for the MTS paradigm, except that at least three days before surgery rats are required to perform at least 3-6 trials of the NMTS paradigm. Rats are then taken off food restriction and undergo surgery. After one week of post-surgery recovery, the food restriction is resumed and once rats are at 85-90% of their post-surgery weight. Finally, rats run 6-15 trials per day until the average performance is below 50% for at least three days in a row.

#### Sequence memory paradigm

Prior to surgery, rats are trained to run back and forth on a linear track (130 cm) for liquid reward. Animals perform two 15 min sessions per day, with a 15-20 min intervening rest in the sleep box. Training continues until rats collected at least 50 rewards in 15 min. Food restriction is then stopped to prepare for surgery. After one week of post-surgery recovery, food restriction is resumed, and rats are put back on the linear track until they perform at least 70 crossings in 15 min. Next, rats are trained on a continuous spatial alternation task in a three arm radial maze following a single-day acquisition procedure that consist of eight 15 min learning sessions. Training on the alternation task is repeated on three days (separated by a rest day), and each day a different arm is chosen as the home arm. On the rest days, the rats perform two 15 min sessions with the same home arm as the day before. Finally, rats are introduced to the sequence memory paradigm as described above until their sequence performance is above 80% by the end of the session, for three days in a row. We found that pretraining rats on the alternation task improved the acquisition of the sequence task.

### Surgical procedure

A custom-designed 3D-printed micro-drive array (Kloosterman et al. (2009)), carrying up to 8 tetrodes (four twisted 0.012 mm polyimide-insulated nickel-chrome wires, angled cut; Sandvik, Kista, Sweden) and 3 stimulation electrodes (two twisted 0.06 mm polyimide-insulated stainless-steel wires, angled cut; California Fine Wire, Grover Beach, CA), was surgically attached to the rat skull using standard aseptic techniques. To induce anesthesia the rat was placed in an induction chamber filled with oxygen (0.5-1 L/m) and 5% isoflurane. Next, its head was securely mounted in a stereotaxic frame after shaving the head, and eyes were protected from drying out and prolonged light exposure with eye ointment and aluminum foil. During the surgery, anesthesia was maintained by administration of 0.5-2% isoflurane through a nose mask and adjusted if necessary, based on vital signs: blood oxygen level, heart rate and breathing rate. The body temperature (measured with a rectal probe) was kept constant using a heating pad. After disinfection of the skin with iodine and ethanol, an incision with a scalpel along the mid-line exposed the skull. Small scratches were carved in the bone plates to allow better adhesion of the dental cement used to fix the implant to the skull. Nine anchoring bone screws were inserted (3 frontal, 2 left parietal, 2 occipital and 1 right parietal) and gaps between the screws and the skull were filled up with bio compatible glue (VetBond) and after extra disinfection (10 min Baytril submersion) all screws were fixed together with dental cement. Next, two craniotomies were drilled, and the dura was removed to allow access to the brain above the HC and ventral hippocampal commissure (VHC) (HC-craniotomy center coordinates: 4 mm posterior to Bregma, 2.5 mm right from the midline; VHC-craniotomy center coordinates: 1.3 mm posterior to Bregma, 0.9 mm right from the midline). Before mounting of the implant, craniotomies were covered in silicon grease to seal the gaps between the edge of the craniotomies and the tetrode-carrying tubes. Additionally mineral oil was applied to the tip of the cannulas prior to implantation which prevents back-filling (and possible clogging) of the tetrode-carrying tubes with cerebrospinal fluid or blood. The implant was fixed to the skull with light-curable dental cement (SDI, Bayswater, Australia) and one screw was wired to the electrode interface board to serve as electrical ground. The skin was sutured in the front and back of the implant with surgical threads (4-0 Silk wax coated braided silk, Sofsilk). While the rat was still under light anesthesia, all tetrodes and stimulation wires were lowered 1 mm into the cortex. Finally, 0.7 ml of saline (anti-dehydration) and 0.3 ml Metacam (anti-inflammatory and pain relieve) were administered subcutaneously. The Metacam injection was repeated in the three days following completion of the surgery.

### Electrophysiological recordings

After approximately one week of post-surgical recovery, 4-5 tetrodes were positioned in the CA1 pyramidal cell layer of the dorsal HC over the course of 3 days to minimize tissue damage. One tetrode was lowered in the white matter above the CA1 cell layer to serve as a reference and one tetrode was placed in the cortex to aid the online ripple detection. Wide-band (0.1 - 4 kHz) signals were sampled at 4 kHz, digitized using a 128-channel data acquisition system (Digilynx SX acquisition system with HS-36 analog headstage and Cheetah software; Neuralynx, Bozeman, MO) and saved to an hard disk for offline analysis.

### Online ripple detection and disruption

A live network stream of digitized multi-channel samples from the Digilynx acquisition system was fed into a quad-core workstation that runs the real-time detection software Falcon (Ciliberti and Kloosterman 2017). Falcon initiates TTL pulses via a microcontrollor board (Arduino UNO) that is connected to a constant-current stimulator (MultiChannel System, Reutlingen, Germany) which generates a biphasic current for electrical stimulation of the VHC. Here, raw signals of 2-4 electrodes (with the clearest ripples) are first filtered in the ripple band (135-255 Hz) using a Chebyshev type-II IIR filter (order 20). Using the neuralynx acquisition system the total round trip latency was below 1 ms (Ciliberti and Kloosterman 2017). After summing of the ripple power (RP) of the different electrodes, ripples were identified based on the following criteria.

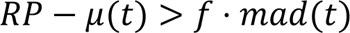

In this equation μ(t) and mad(t) are the running estimates of the mean and mean absolute deviation calculated on the ripple power, and f is a multiplier set to a value in the range 5-13. The value of f was adjusted for each daily session to maximize the ratio of positive over false ripple detections. The RP is the root mean square of the filtered signal. Estimates of μ(t) and mad(t) were computed using an exponential moving average filter with span set to 7 seconds. Estimates were not updated during a 50 ms window after each detection minimize the contribution of the ripple. to avoid multiple detections of the same event. To avoid false detections due to heavy head movements or chewing artifacts, the same ripple detection procedure was applied to signals from the cortical electrode. As ripples are not present in the cortex but artifacts like the ones mentioned above are, detections were marked as false positive if a cortical detection occurred within a fixed time window ([-40, 1.5] ms) around a HC detection. All not positively marked detections triggered the generation of a second TTL pulse by Arduino UNO either immediately (disruption condition) or after a random 150-250 ms delay between 150-250 ms. This TTL-pulse was sent to a constant-current stimulator (MultiChannel System, Reutlingen, Germany) to electrically stimulate the VHC. Both the detection event and the time of stimulation (= stimulation event) were sent to the Neuralynx acquisition system for logging. The amplitude of the biphasic electrical pulses (0.2 ms duration) varied from 50-250 A and was set in each session to the lowest amplitude that resulted in consistent disruption of hippocampal ripple events. To avoid over-stimulation that could lead to damage of the surrounding tissue, no stimulations could occur within a 150 ms lockout period after a prior stimulation. In the disruption condition this is achieved by discarding detections in a 150ms window after each detection. In the stimulated control condition, all detections that would trigger a stimulation that falls withing a 150ms window after the delayed stimulation are were discarded. Ripple detection and disruption started after rats acquired the task rules reached according to the learning criteria specific for the behavioral task. In the NMTS paradigm ripple detection and/or disruption only occurs only during a trial (i.e., both during and in between the instruction and test phase), but not in the inter-trial time. For every trial one of the control/disruption conditions was picked pseudo-randomly. In the MTS paradigm ripple detection and/or disruption was applied during both the trials and the inter-trial times intervals. In the sequence memory paradigm, ripple detection and/or disruption was applied during both run epochs and rest epoch. In both the MTS and sequence memory paradigm the stimulation condition was constant through the whole session.

### Quantification and statistical analysis

Analysis of neural and behavioral data was performed using Python and its scientific extension modules (Millman and Aivazis 2011), augmented with custom Python and C++ toolboxes. Throughout the result section, all description summary statistics are in the format ‘mean [99% CI],’ unless stated otherwise.

### Behavior

Position and speed of the rat were tracked using LED lights mounted on one of the headstages and an overhead video camera (25 Hz). Entry into a maze arm was detected as soon as the animal moved past the first 20 cm of the arm (regardless of the rat actually reaching the reward platform). To assess stereotypical behavior in the MTS task, we defined for every session a stereotypy index. First, a string representing the sequence of arm visits in a trial was created for each of the last 10 trials in a session (for example, ‘1345’ or ‘5471’). The stereotypy index was defined as the average similarity (based on the normalized Levenshtein distance) between the strings of all trial pairs. The Levenshtein distance measures the minimum number of single character edits that are necessary to change one string into another, and for two strings a and b with lengths |a| and |b| is computed according to the following formula:

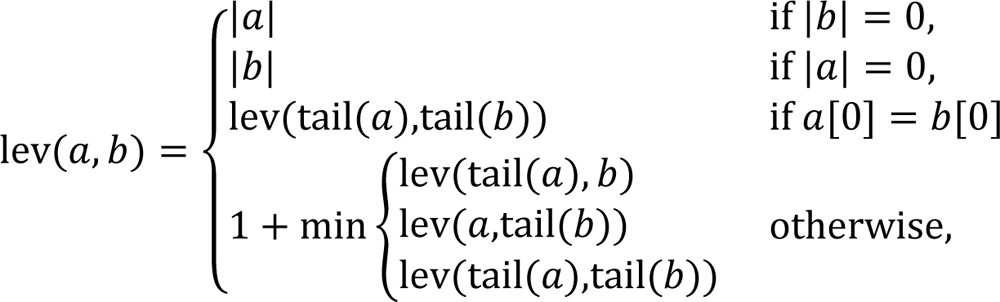

with tail(x) equal to all of string x except for the first character. The similarity is then computed as one minus the normalized Levenshtein distance:

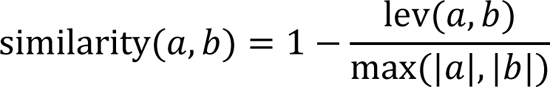

The shuffled distributions used to compute the z-scores for the circularity and stereotypy quantifications of every MTS session, were created by by randomizing the visit order of each trial 2000 times.

For behavioral analysis of the sequence paradigm, only sessions in which the rat performed at least 150 visits in the first run epoch are considered. After this selection, data from run 1 and run 2 are merged and we only consider the first 400 visits to assess the (learning) behavior. To compute smoothed learning curves we use a Gaussian kernel with a bandwidth of 50 trials. Next, we calculated the corresponding binomial probability that the (smoothed) number of correct responses is significantly higher (alpha = 0.01) than would be expected by chance (0.5) and define the last trial at which this criteria is met. If the criteria are never met, no learning visit was defined.

### Offline ripple detection

Ripple events were detected in the hippocampal field potential for all trials and sessions of the control and stimulated control conditions. For the stimulated control condition, the stimulation artifact is first removed and replaced by cubic interpolation of the signal in a 10ms window around the stimulation. To detect ripples offline, the local field potential recorded from 1-3 tetrodes was filtered in the ripple frequency band (140-225 Hz). The ripple envelope was computed as the absolute value of the Hilbert-transformed ripple signal, averaged across the recording sites and smoothed with a Gaussian kernel (bandwidth 15 ms). Finally, ripple events were detected when the ripple envelope exceeded a high threshold of med + 9 ⋅ (Q3 − med). Here, med and Q3 represent the median and upper quartile range of the envelope. To determine the start and end time of each ripple, as second the threshold was computed with the same method as before, but now only using the signal statistics of a 30ms window before each detected ripple (−100ms to −70ms) and a multiplier of 0.4 instead of 9. Ripple events that were separated by less than 20 ms were merged into one, and events with a duration shorter than 40 ms were excluded. Offline ripple detection was performed on the same task epochs when online ripple detection was performed. It should be noted that the multiplier for the percentile-based threshold, that we use, should not be compared directly to the multiplier for z-score based threshold that is used in previous studies because of the different statistics being used (median / upper quartile range vs mean / standard deviation). When computing the equivalent z-score multiplier that produces the same absolute threshold as our percentile multiplier, we find that these multiplier values are comparable to the 3-6 standard deviations reported in previous papers (Girardeau et al. 2009; Jadhav et al. 2012; Fernández-Ruiz et al. 2019) (Figure 4–figure supplement 1d,e).

### Evaluation of online ripple detection and disruption

Online ripple detection accuracy was verified for all trials and sessions of the control and stimulated control conditions. Online detected ripples were compared to offline detected ripples to identify the fraction of offline ripples that were also detected online (true positive rate or TPR) and the fraction of online detections that did not correspond to an offline detected ripples (false discovery rate, FDR). To assess the disruption of ripples after a closed-loop stimulation of the VHC, the mean ripple envelope after each detection/stimulation was computed in a 30 ms time window (from +20 ms to +50 ms) and normalized to the mean ripple envelope in a 30 ms time window (from −50ms to −20ms) before detection/stimulation. The ripple envelope was computed as described above, including the removal of the stimulation artifact for the disruption session. The artifact removal was also performed at the time of detection for the stimulated control sessions to have a fair comparison between the disruption and stimulated control sessions.

### Statistics

To test a difference in means between two (unpaired) samples, we used the Mann-Whitney test. To test the difference in means between three (unpaired) samples we used the Kruskal-Wallis test.

### Data and software availability

Falcon software for closed-loop ripple detection and code for analysis are publicly available at http://www.bitbucket.org/kloostermannerflab. Source data are deposited in the following Open Science Framework project repository: https://osf.io/anc4m.

## Supplementary Figures

**Figure 1–figure supplement 1:**
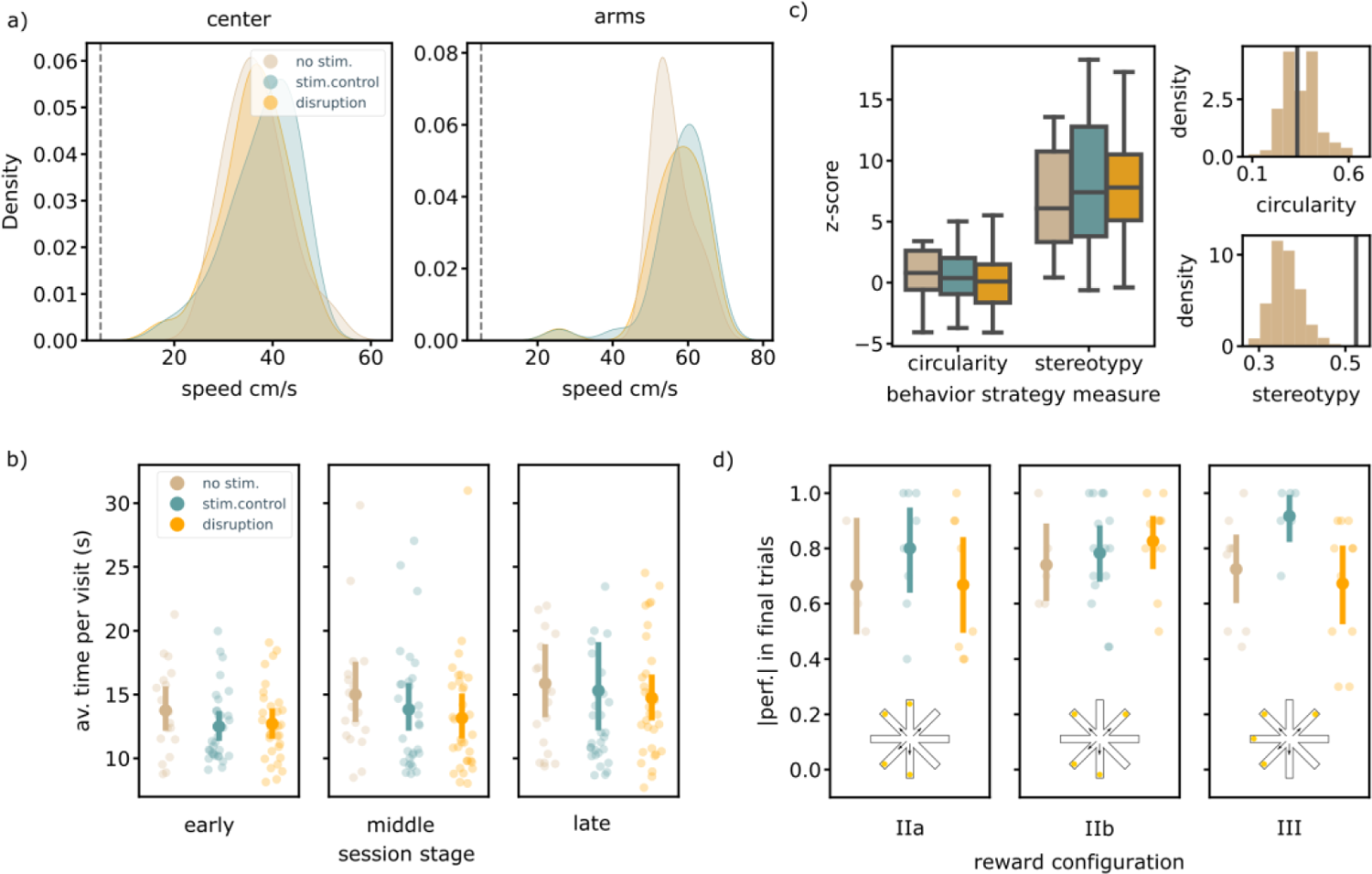
Additional behavioral analyses MTS task show no effect of ripple disruption. a) Speed distributions of the average speed at the reward platforms, central platforms or arms in a session per stimulation protocol. The dashed line indicates 5cm/s, the threshold used to determine immobility periods. The speed is significantly lower on the central platform vs. the arms in all three stimulation conditions; disruption; center: 36.42 [33.20,39.10]cm/s, arms: 57.15 [51.93,59.65]cm/s, Mann-Whitney test: U=43.00, p= 1.3 × 10^−10^, stim.control; center: 38.06 [34.59,40.76]cm/s, arms: 58.12 [52.94,60.59]cm/s, Mann-Whitney test: U=30.00, p= 1 × 10^−10^, no stim.; center: 36.78 [33.67,41.12]cm/s, arms: 55.95 [53.38,59.50]cm/s, Mann-Whitney test: U=2.00, p= 1 × 10^−6^. The speed on each region is not significantly different across stimulation protocols; Kruskal-Wallis test, center: H=2.03, p=0.36, arms: H=4.14, p=0.13 b) The average time per visit in the early (1-8), middle (9-17) and late (18-25) trials of a session is not significantly different in the disruption vs. control sessions; early; disruption: 12.71 [11.51,14.17]s, stim.control: 12.49 [11.43,14.04]s, no stim.: 13.75 [11.99,15.78]s, Kruskal-Wallis test: H=1.74, p=0.42, middle; disruption: 13.15 [11.57,16.04]s, stim.control: 13.83 [12.02,16.43]s, no stim.: 15.00 [12.86,19.31]s, Kruskal-Wallis test: H=2.25, p=0.32, late; disruption: 14.73 [12.80,17.08]s, stim.control: 15.31 [12.47,22.31]s, no stim.: 15.87 [13.07,20.64]s, Kruskal-Wallis test: H=1.52, p=0.47. c) (left) Distributions of the z-scores for the behavioral strategy measures (fraction (counter)clockwise trials and stereotypy index) of each session compiled from shuffled distributions (see methods) The z-scores do not change due to ripple disruption (Kruskal-Wallis test, circularity: H=0.42, p=0.81, stereotypy: H=0.84, p=0.66). (right) Example distributions of each behavioral strategy measure compiled from the shuffled distributions. Solid vertical line indicates the measure for the example session. d) The average performance in the final trials for sessions sorted per arm configuration; average performance in final trials; IIa; disruption: 0.67 [0.46,0.88], stim.control: 0.80 [0.56,0.94], no stim.: 0.67 [0.50,0.90], Kruskal-Wallis test: H=1.58, p=0.45, IIb; disruption: 0.83 [0.66,0.91], stim.control: 0.78 [0.65,0.89], no stim.: 0.74 [0.60,0.94], Kruskal-Wallis test: H=1.13, p=0.57, III; disruption: 0.67 [0.45,0.81], stim.control: 0.92 [0.73,0.98], no stim.: 0.72 [0.55,0.87], Kruskal-Wallis test: H=6.21, p=0.045, average number of visits in final trials; IIa; disruption: 4.73 [4.26,5.48], stim.control: 4.26 [4.06,4.62], no stim.: 4.77 [4.40,5.40], Kruskal-Wallis test: H=4.08, p=0.13, IIb; disruption: 4.26 [4.11,4.49], stim.control: 4.41 [4.19,4.73], no stim.: 4.38 [4.08,4.80], Kruskal-Wallis test: H=0.87, p=0.65, III; disruption: 4.42 [4.22,4.67], stim.control: 4.10 [4.00,4.23], no stim.: 4.40 [4.17,4.67], H=6.07, p=0.048, learning trial; IIa; disruption: 10.38 [7.21,14.82], stim.control: 10.00 [7.38,16.00], no stim.: 14.00 [6.00,25.00], Kruskal-Wallis test: H=0.40, p=0.82, IIb; disruption: 12.18 [8.18,18.00], stim.control: 9.00 [7.55,10.00], no stim.: 8.80 [5.80,12.00], Kruskal-Wallis test: H=0.71, p=0.7, III; disruption: 10.27 [7.36,16.61], stim.control: 9.83 [6.33,14.67], no stim.: 9.00 [6.44,11.56], Kruskal-Wallis test: H=0.07, p=0.96.

**Figure 2–figure supplement 1:**
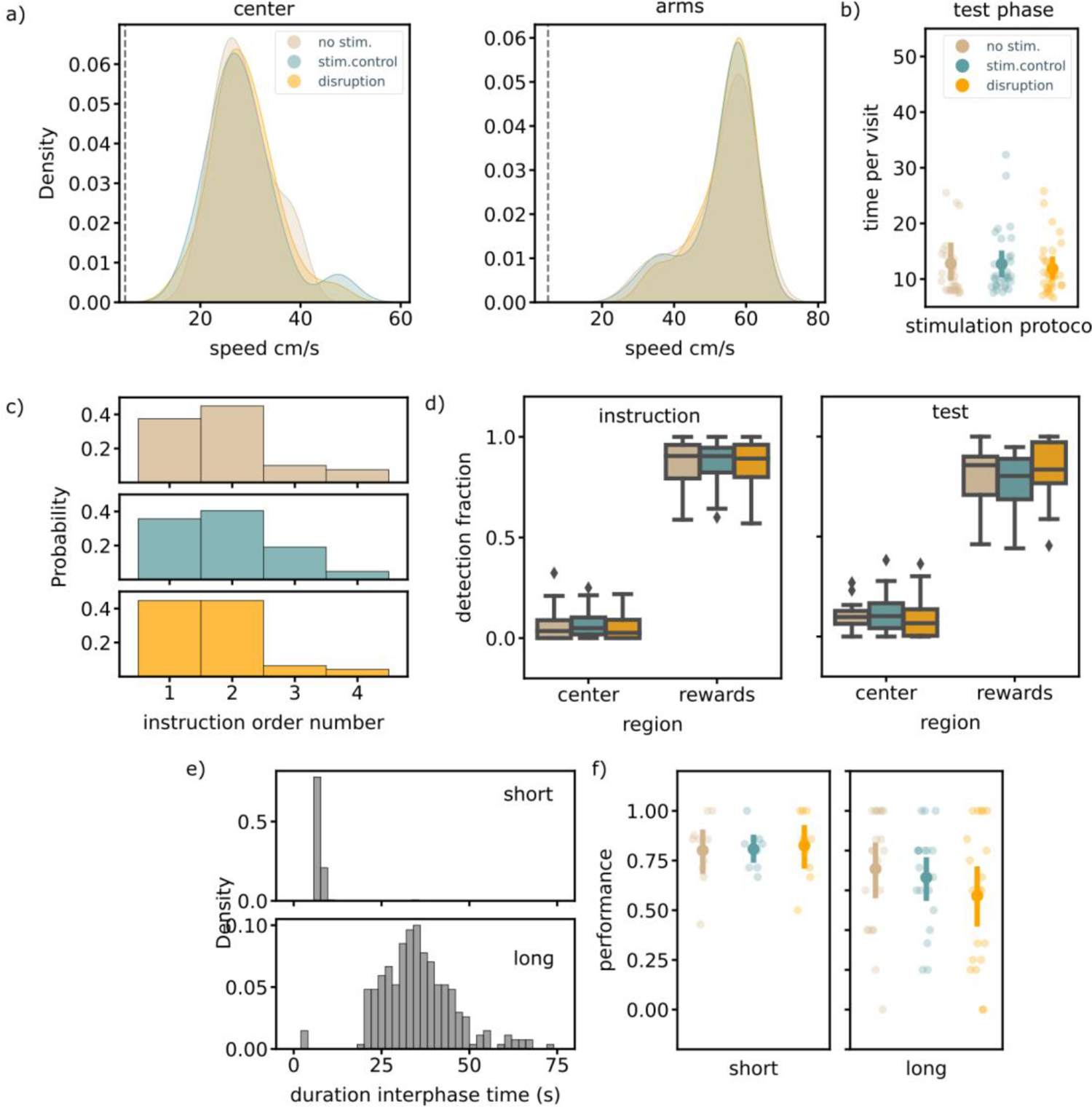
Additional behavioral analyses NMTS task show no effect of ripple disruption. a) Speed distributions of the average speed at the reward platforms, central platforms or arms in a session per stimulation protocol. The dashed line indicates 5cm/s, the threshold used to determine immobility periods: disruption; arms: 54.02 [49.42,57.02]cm/s, center: 28.66 [26.06,31.96]cm/s, stim.control; arms: 53.24 [48.08,56.52]cm/s, center: 28.39 [25.83,32.37]cm/s, no stim.; arms: 53.09 [47.96,56.57]cm/s, center: 28.67 [26.31,31.49]cm/s. There is no significant difference due to ripple disruption; Kruskal-Wallis test; arms: H=0.09, p=0.96, center: H=0.42, p=0.81. b) The average time per visit in the test phase of a trial per stimulation protocol; disruption: 11.24 [10.32,12.49]s, stim.control: 11.67 [10.68,12.98]s, no stim.: 10.36 [9.62,11.93]s, Kruskal-Wallis test: H=6.06, p=0.048. c) The distribution of the instruction phase order number of the wrong visit in incorrect trials, for all three stimulation protocols. d) The fraction of ripple detections during the instruction and test phase on the rewards and central platforms; instruction: rewards; disruption: 0.86 [0.79,0.91], stim.control: 0.87 [0.81,0.91], no stim.: 0.88 [0.82,0.92], center; disruption: 0.05 [0.03,0.09], stim.control: 0.07 [0.05,0.11], no stim.: 0.06 [0.03,0.11], arms; disruption: 0.06 [0.04,0.11], stim.control: 0.05 [0.03,0.08], no stim.: 0.05 [0.03,0.08], test: rewards; disruption: 0.85 [0.77,0.90], stim.control: 0.78 [0.72,0.84], no stim.: 0.80 [0.74,0.86], center; disruption: 0.09 [0.06,0.15], stim.control: 0.12 [0.08,0.17], no stim.: 0.10 [0.08,0.14], arms; disruption: 0.05 [0.03,0.08], stim.control: 0.08 [0.05,0.12], no stim.: 0.09 [0.05,0.14]). The reward bias is not significantly different between stimulation protocols: Kruskal-Wallis test, instruction: H=0.43, p=0.8, test: H=4.02, p=0.13. e) The distributions for the time between instruction and test phase; long: 35.38 [33.76,37.03]s, short; 7.29 [7.06,8.10]s, Mann-Whitney test: U=854.00, p=6.7 × 10^−68^. f) The average trial performance per stimulation protocol in sessions with a short or long interphase delay; long; disruption: 0.60 [0.46,0.72], stim.control: 0.67 [0.52,0.77], no stim.: 0.68 [0.53,0.79], H=1.59, p=0.45, short; disruption: 0.82 [0.62,0.91], stim.control: 0.82 [0.63,0.92], no stim.: 0.81 [0.64,0.90], H=0.04, p=0.98. Statistics for the number of correct visits (no figure shown): long; disruption: 3.39 [3.08,3.59], stim.control: 3.59 [3.39,3.73], no stim.: 3.49 [3.18,3.68], H=1.81, p=0.4, short; disruption: 3.73 [3.29,3.89], stim.control: 3.75 [3.32,3.88], no stim.: 3.67 [3.27,3.85], H=0.06, p=0.97.

**Figure 3–figure supplement 1:**
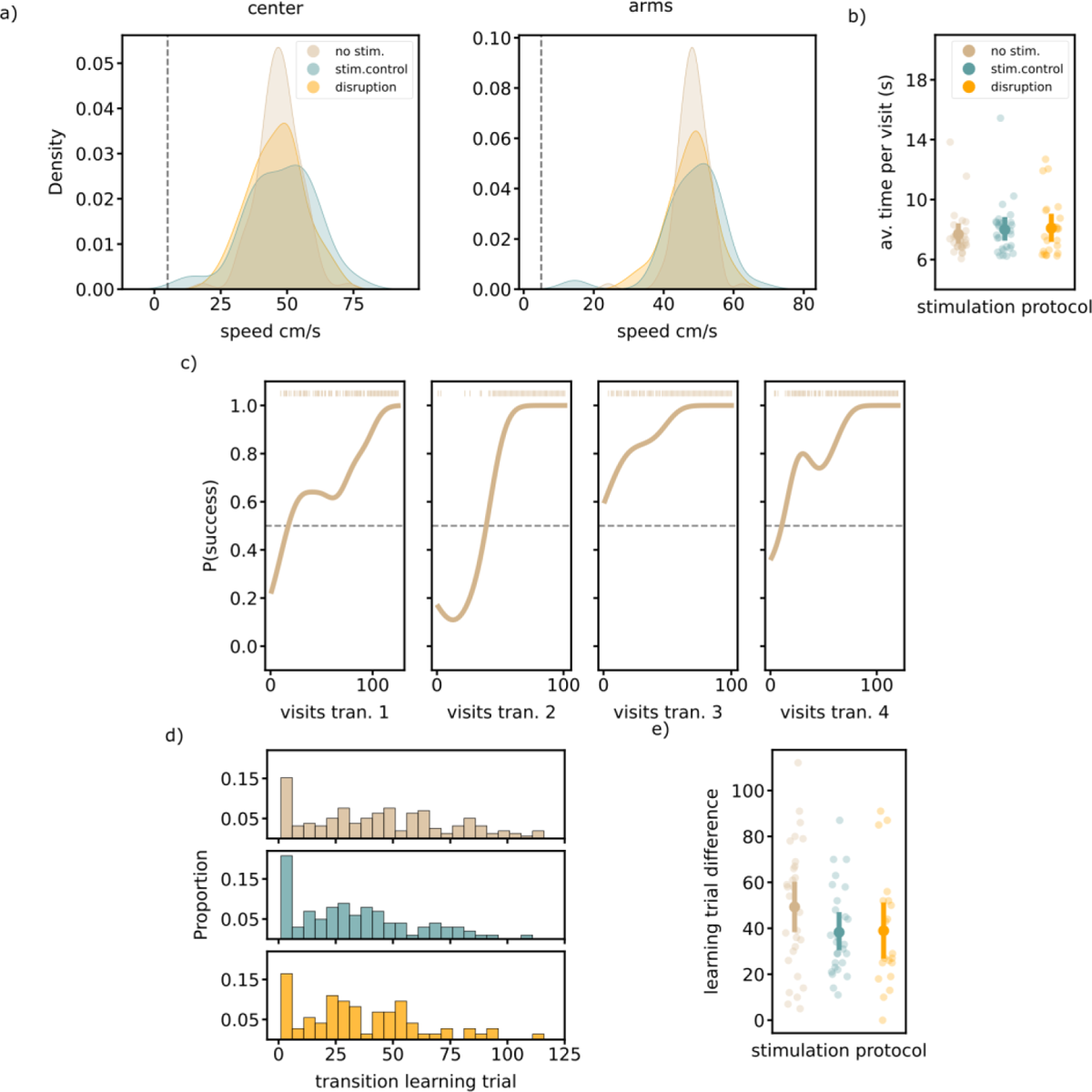
Additional behavioral analyses SEQ task show no effect of ripple disruption. a) Speed distributions of the average speed at the reward platforms, central platforms or arms in a session per stimulation protocol. The dashed line indicates 5cm/s, the threshold used to determine immobility periods; center; disruption: 46.01 [42.75,49.49]cm/s, stim. control 47.56 [43.05,51.80]cm/s, no stim.: 46.93 [44.85,48.84]cm/s, Kruskal-Wallis test: H=1.20, p=0.55, arms; disruption: 47.49 [45.27,49.49]cm/s, stim.control: 48.43 [44.79,50.91]cm/s, no stim.: 48.29 [46.96,49.31]cm/s, Kruskal-Wallis test: H=1.79, p=0.41. b) The average time per visit in the run 1 and run 2 of a session per stimulation protocol; disruption: 7.72 [7.67,7.78]s, stim.control: 7.73 [7.68,7.78]s, no stim.: 7.50 [7.46,7.53]s, Kruskal-Wallis test: H=0.77, p=0.68. c) Learning curves for individual transitions in one example session. The dashed line indicates the learning criteria, the brown lines on top indicate correct visits. d) Distribution of transition learning trials for sessions in each stimulation protocol; Kruskal-Wallis test: H=10.62, p=0.0049, post-hoc tests; disruption - stim.control: U=3946.00, p=0.43, disruption - no stim.: U=4801.50, p=0.041, stim.control - no stim.: U=9782.50, p=0.0021. e) The learning trial difference for each session, defined by the difference between the learning trial of the slowest and fasted learned transition; disruption: 38.95 [26.30,55.04], stim.control: 38.30 [30.13,49.11], no stim.: 49.31 [36.16,62.20], Kruskal-Wallis test: H=3.10, p=0.21. Only sessions in which at least two transitions are learned are included. (disruption: N= 20, stim.control: N= 27, no stim.: N= 29)

**Figure 4–figure supplement 1:**
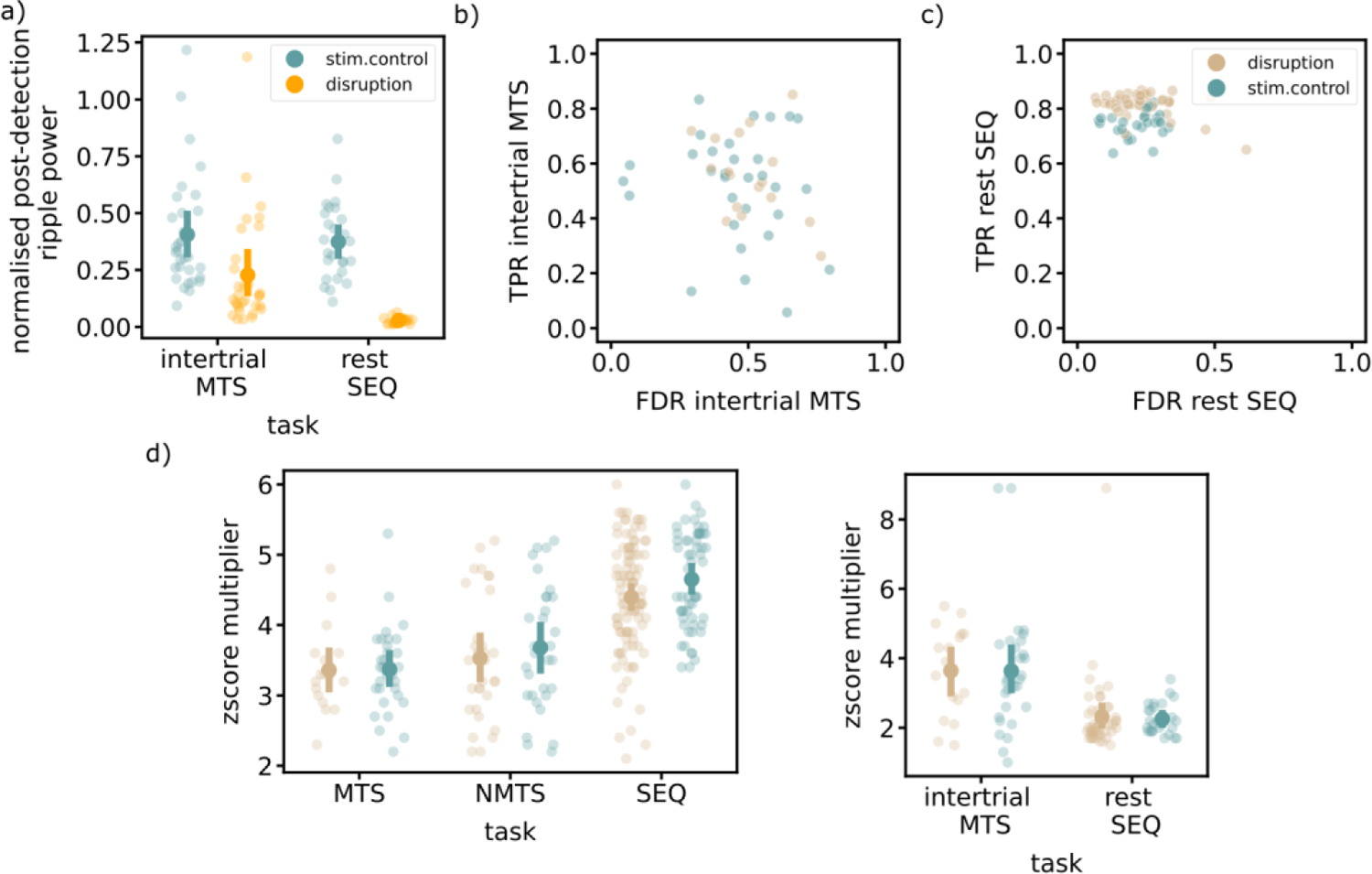
Ripple detection and disruption quantifications in rest epochs. a) Normalized post-detection ripple power for all tree tasks in rest epochs; inter-trial times (in center of the maze) for MTS task, rest epochs (in sleep box) for SEQ task. One dot represents the average over one session; Mann-Whitney test, MTS: U=754.00, p=6.1 × 10^−5^, SEQ: U=621.00, p= 8 × 10^−10^. b) and c) True positive rate versus False discovery rate for inter-trial times in MTS task and rest epochs in SEQ task respectively. One dot represents the average over on session. d) The quivalent z-score multiplier that produces the same absolute ripple threshold as our percentile multiplier for all run epochs (left) and rest epochs (right) for all three tasks. One dot represents the average over one session; RUN: MTS: stim.control, 3.37 [3.12,3.70], no stim., 3.36 [3.05,3.81], NMTS: stim.control, 3.68 [3.29,4.08], no stim., 3.53 [3.14,3.96], SEQ: stim.control 4.65 [4.42,4.87], no stim. 4.40 [4.17,4.60], REST: MTS: stim. control 3.64 [3.02,4.64], no stim. 3.64 [2.81,4.40], SEQ: stim. control2.26 [2.07,2.51], no stim.2.31 [2.09,3.17]

